## Supplementary material for "Non-invasive prenatal testing by low coverage genomic sequencing: Detection limits of screened chromosomal microdeletions"

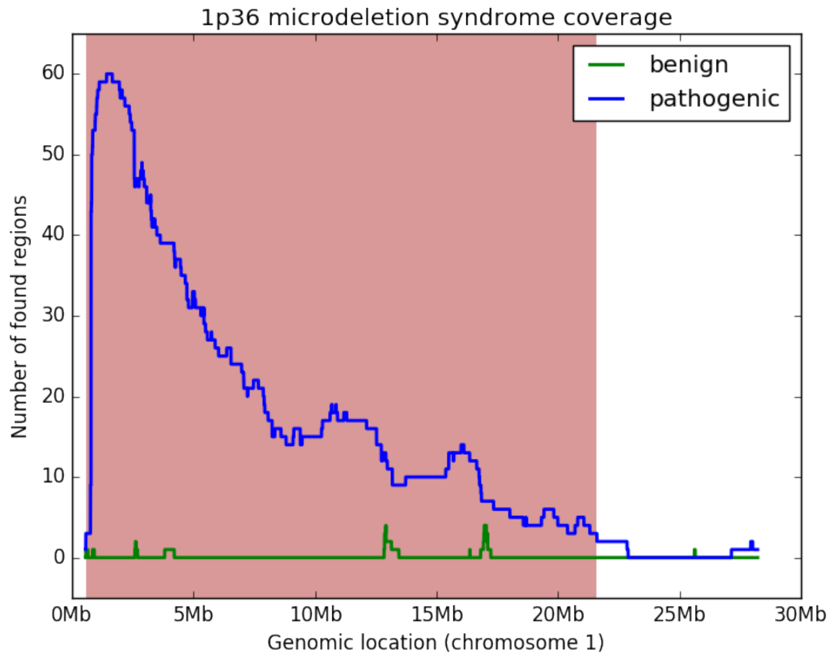


***Supplementary Figure 1.:*** *Coverage plot representing genomic position of critical region of 1p36 microdeletion syndrome. Coverage of pathogenic and likely pathogenic deletions is denoted by blue, while coverage of benign and likely benign microdeletions is in green. The critical region is visualized in red.*

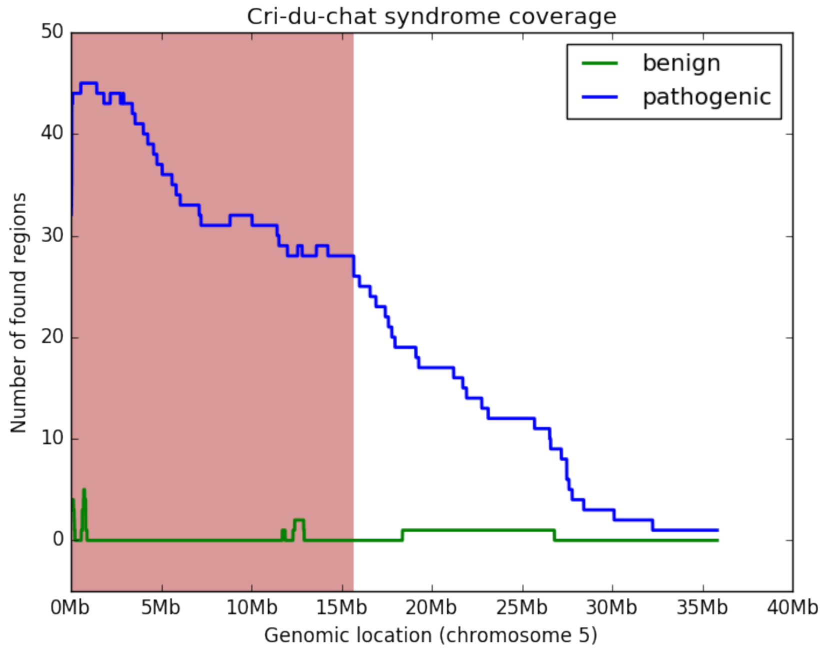


***Supplementary Figure 2.:*** *Coverage plot representing genomic position of critical region of Cri-Du-Chat syndrome. Coverage of pathogenic and likely pathogenic deletions is denoted by blue, while coverage of benign and likely benign microdeletions is in green. The critical region is visualized in red.*

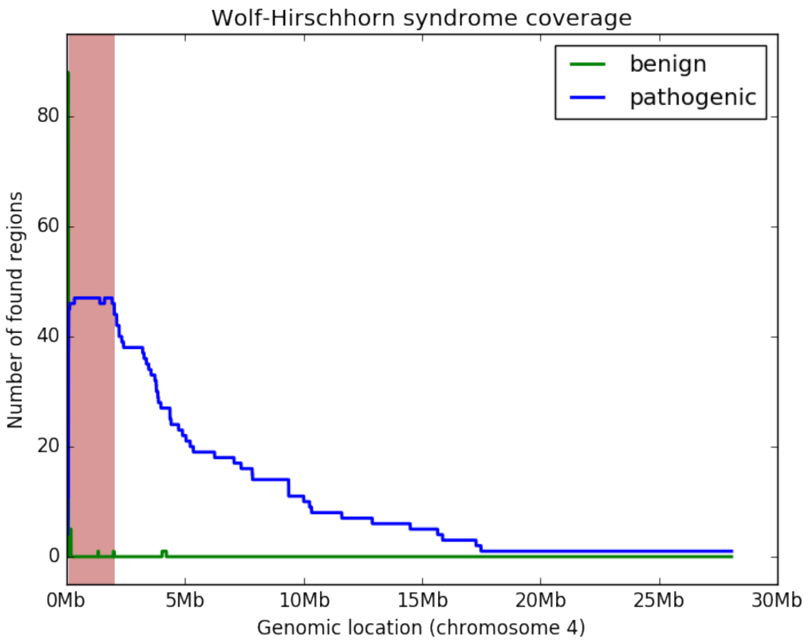


***Supplementary Figure 3.:*** *Coverage plot representing genomic position of critical region of Wolf-Hirschhorn syndrome. Coverage of pathogenic and likely pathogenic deletions is denoted by blue, while coverage of benign and likely benign microdeletions is in green. The critical region is visualized in red.*

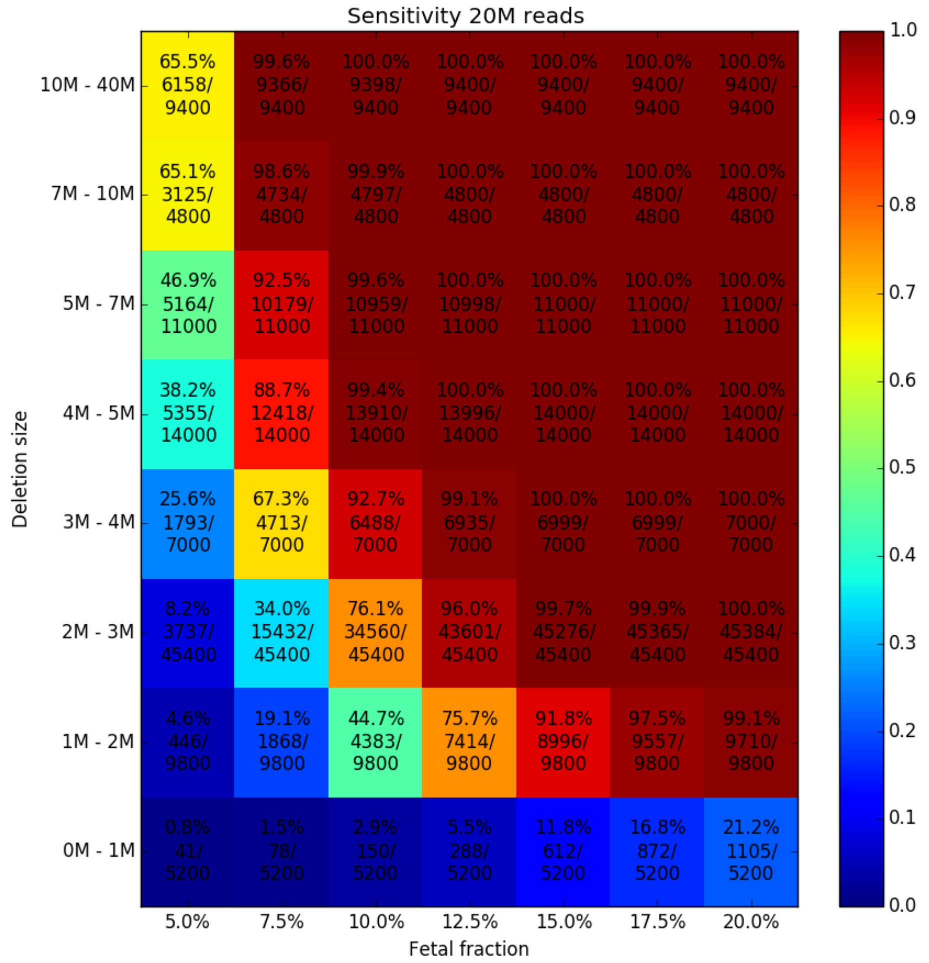


***Supplementary Figure 4.:*** *Sensitivity of the prediction for different fetal fraction and microdeletion size. Read count was set to 20M in each sample. Detections 2Mb away from critical region are reported.*

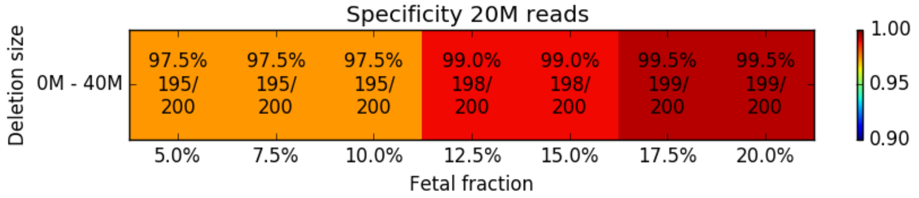


***Supplementary Figure 5.:*** *Specificity of the prediction for different fetal fraction and microdeletion size. Read count was set to 20M in each sample. Detections 2Mb away from critical region are reported.*

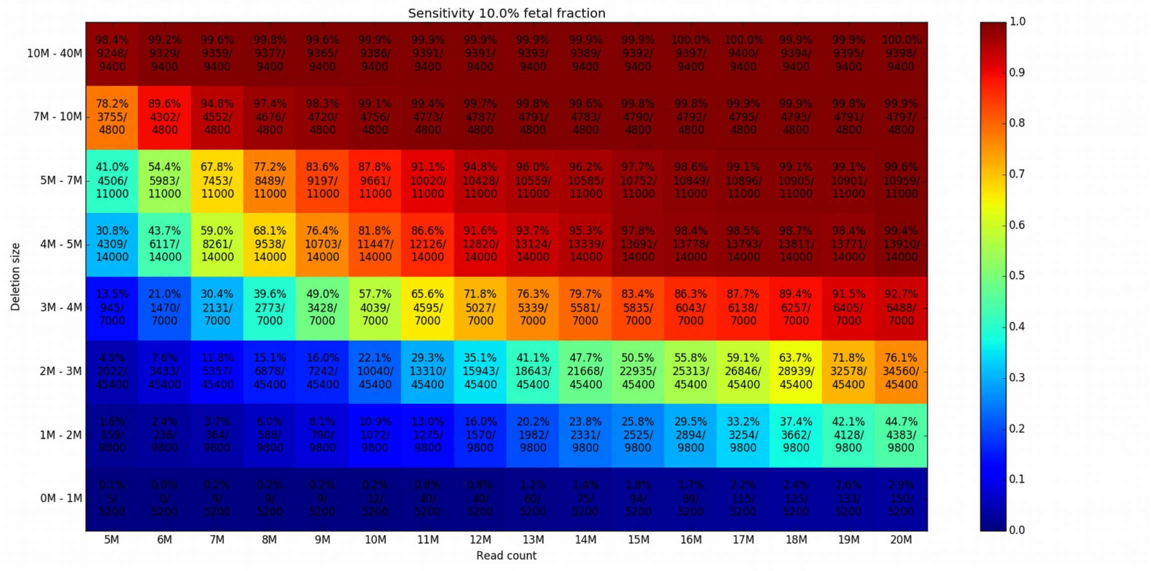


***Supplementary Figure 6.:*** *Sensitivity of the prediction for different read count and microdeletion size. Fetal fraction was set to 10% for all samples. Detections 2Mb away from critical region are reported.*

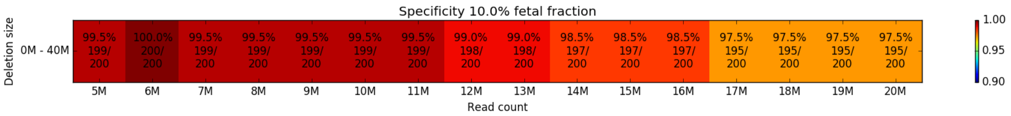


***Supplementary Figure 7.:*** *Specificity of the prediction for different read count and microdeletion size. Fetal fraction was set to 10% for all samples. Detections 2Mb away from critical region are reported.*

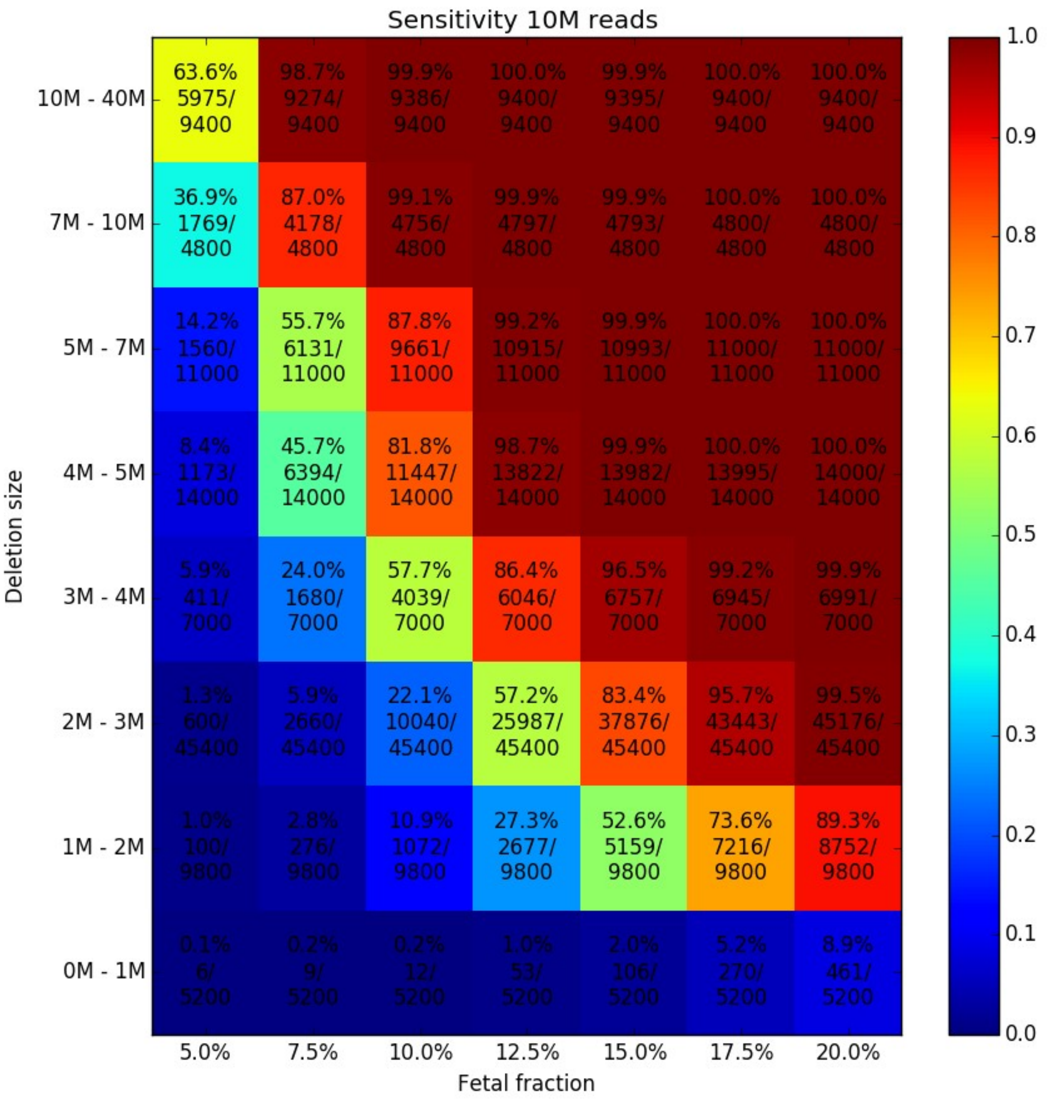


***Supplementary Figure 8.:*** *Sensitivity of the prediction for different fetal fraction and microdeletion size. Read count was set to 10M in each sample. Detections 2Mb away from critical region are reported.*

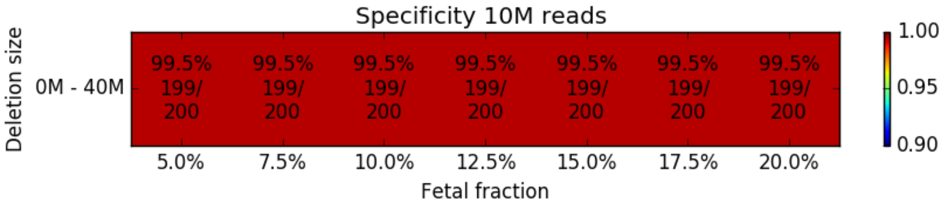


***Supplementary Figure 9.:*** *Specificity of the prediction for different fetal fraction and microdeletion size. Read count was set to 10M in each sample. Detections 2Mb away from critical region are reported.*

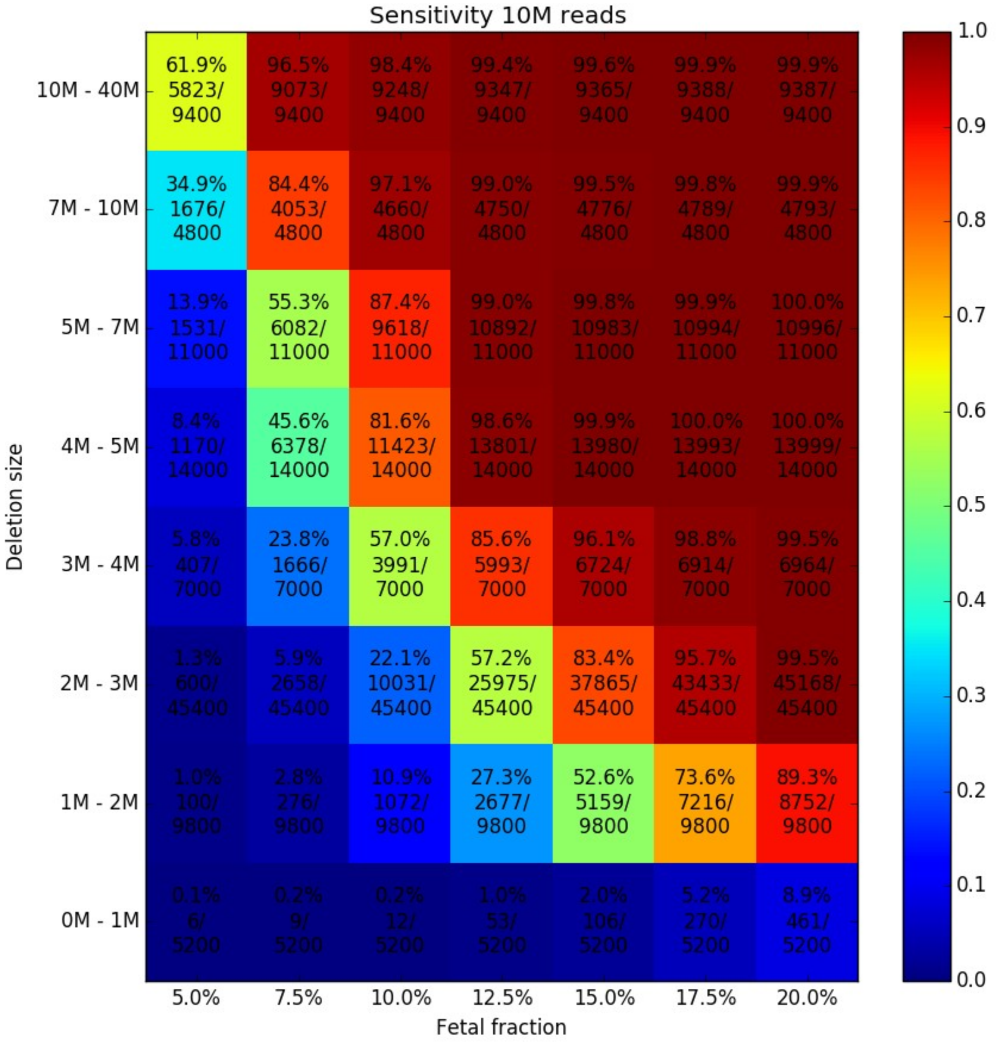


***Supplementary Figure 10.:*** *Sensitivity of the prediction for different fetal fraction and microdeletion size. Read count was set to 10M in each sample. Detections 2Mb away from critical region are NOT reported.*

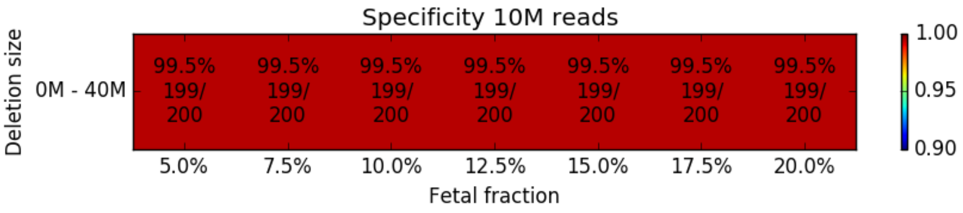


***Supplementary Figure 11.:*** *Specificity of the prediction for different fetal fraction and microdeletion size. Read count was set to 10M in each sample. Detections 2Mb away from critical region are NOT reported.*

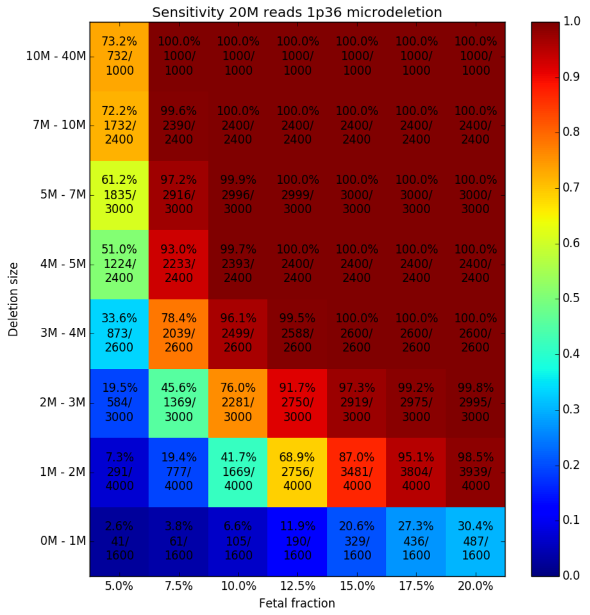
Sensitivity 20M reads 1p36 microdeletion
Deletion size
Fetal fraction
10M - 40M
73.2% 732/ 1000
100.0% 1000/ 1000
100.0% 1000/ 1000
100.0% 1000/ 1000
100.0% 1000/ 1000
100.0% 1000/ 1000
100.0% 1000/ 1000
7M - 10M
72.2% 1732/ 2400
99.6% 2390/ 2400
100.0% 2400/ 2400
100.0% 2400/ 2400
100.0% 2400/ 2400
100.0% 2400/ 2400
100.0% 2400/ 2400
5M - 7M
61.2% 1835/ 3000
97.2% 2916/ 3000
99.9% 2996/ 3000
100.0% 2999/ 3000
100.0% 3000/ 3000
100.0% 3000/ 3000
100.0% 3000/ 3000
4M - 5M
51.0% 1224/ 2400
93.0% 2233/ 2400
99.7% 2393/ 2400
100.0% 2400/ 2400
100.0% 2400/ 2400
100.0% 2400/ 2400
100.0% 2400/ 2400
3M - 4M
33.6% 873/ 2600
78.4% 2039/ 2600
96.1% 2499/ 2600
99.5% 2588/ 2600
100.0% 2600/ 2600
100.0% 2600/ 2600
100.0% 2600/ 2600
2M - 3M
19.5% 584/ 3000
45.6% 1369/ 3000
76.0% 2281/ 3000
91.7% 2750/ 3000
97.3% 2919/ 3000
99.2% 2975/ 3000
99.8% 2995/ 3000
1M - 2M
7.3% 291/ 4000
19.4% 777/ 4000
41.7% 1669/ 4000
68.9% 2756/ 4000
87.0% 3481/ 4000
95.1% 3804/ 4000
98.5% 3939/ 4000
0M - 1M
2.6% 41/ 1600
3.8% 61/ 1600
6.6% 105/ 1600
11.9% 190/ 1600
20.6% 329/ 1600
27.3% 436/ 1600
30.4% 487/ 1600
5.0% 7.5% 10.0% 12.5% 15.0% 17.5% 20.0%
0.0 0.1 0.2 0.3 0.4 0.5 0.6 0.7 0.8 0.9 1.0

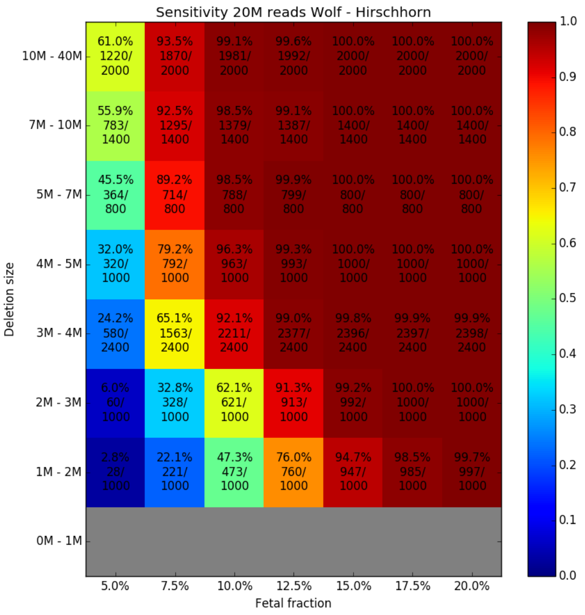
Sensitivity 20M reads Wolf - Hirschhorn
Deletion size
Fetal fraction
10M - 40M
61.0% 1220/ 2000
93.5% 1870/ 2000
99.1% 1981/ 2000
99.6% 1992/ 2000
100.0% 2000/ 2000
100.0% 2000/ 2000
100.0% 2000/ 2000
7M - 10M
55.9% 783/ 1400
92.5% 1295/ 1400
98.5% 1379/ 1400
99.1% 1387/ 1400
100.0% 1400/ 1400
100.0% 1400/ 1400
100.0% 1400/ 1400
5M - 7M
45.5% 364/ 800
89.2% 714/ 800
98.5% 788/ 800
99.9% 799/ 800
100.0% 800/ 800
100.0% 800/ 800
100.0% 800/ 800
4M - 5M
32.0% 320/ 1000
79.2% 792/ 1000
96.3% 963/ 1000
99.3% 993/ 1000
100.0% 1000/ 1000
100.0% 1000/ 1000
100.0% 1000/ 1000
3M - 4M
24.2% 580/ 2400
65.1% 1563/ 2400
92.1% 2211/ 2400
99.0% 2377/ 2400
99.8% 2396/ 2400
99.9% 2397/ 2400
99.9% 2398/ 2400
2M - 3M
6.0% 60/ 1000
32.8% 328/ 1000
62.1% 621/ 1000
91.3% 913/ 1000
99.2% 992/ 1000
100.0% 1000/ 1000
100.0% 1000/ 1000
1M - 2M
2.8% 28/ 1000
22.1% 221/ 1000
47.3% 473/ 1000
76.0% 760/ 1000
94.7% 947/ 1000
98.5% 985/ 1000
99.7% 997/ 1000
0M - 1M
5.0% 7.5% 10.0% 12.5% 15.0% 17.5% 20.0%
0.0 0.1 0.2 0.3 0.4 0.5 0.6 0.7 0.8 0.9 1.0

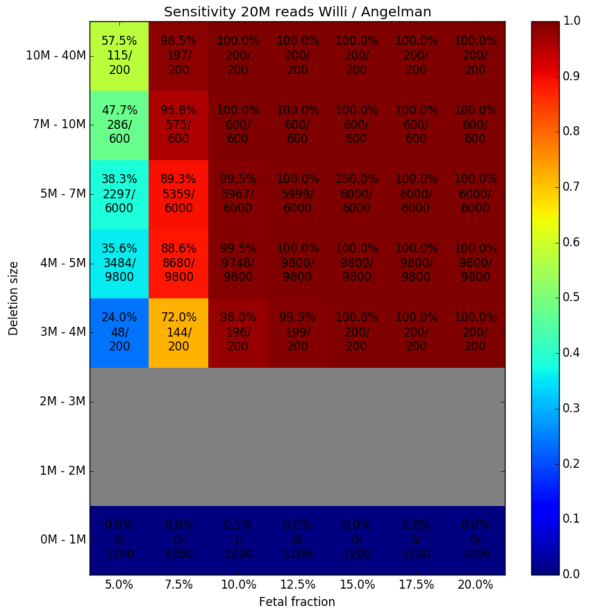
Sensitivity 20M reads Willi / Angelman
Deletion size
Fetal fraction
10M - 40M
57.5% 115/ 200
98.5% 197/ 200
100.0% 200/ 200
100.0% 200/ 200
100.0% 200/ 200
100.0% 200/ 200
100.0% 200/ 200
7M - 10M
47.7% 286/ 600
95.8% 575/ 600
100.0% 600/ 600
100.0% 600/ 600
100.0% 600/ 600
100.0% 600/ 600
100.0% 600/ 600
5M - 7M
38.3% 2297/ 6000
89.3% 5359/ 6000
99.5% 5967/ 6000
100.0% 5999/ 6000
100.0% 6000/ 6000
100.0% 6000/ 6000
100.0% 6000/ 6000
4M - 5M
35.6% 3484/ 9800
88.6% 8680/ 9800
99.5% 9748/ 9800
100.0% 9800/ 9800
100.0% 9800/ 9800
100.0% 9800/ 9800
100.0% 9800/ 9800
3M - 4M
24.0% 48/ 200
72.0% 144/ 200
98.0% 196/ 200
99.5% 199/ 200
100.0% 200/ 200
100.0% 200/ 200
100.0% 200/ 200
2M - 3M
1M - 2M
0M - 1M
0.0% 0/ 1200
0.0% 0/ 1200
0.1% 1/ 1200
0.0% 0/ 1200
0.0% 0/ 1200
0.0% 0/ 1200
0.0% 0/ 1200
5.0% 7.5% 10.0% 12.5% 15.0% 17.5% 20.0%
0.0 0.1 0.2 0.3 0.4 0.5 0.6 0.7 0.8 0.9 1.0

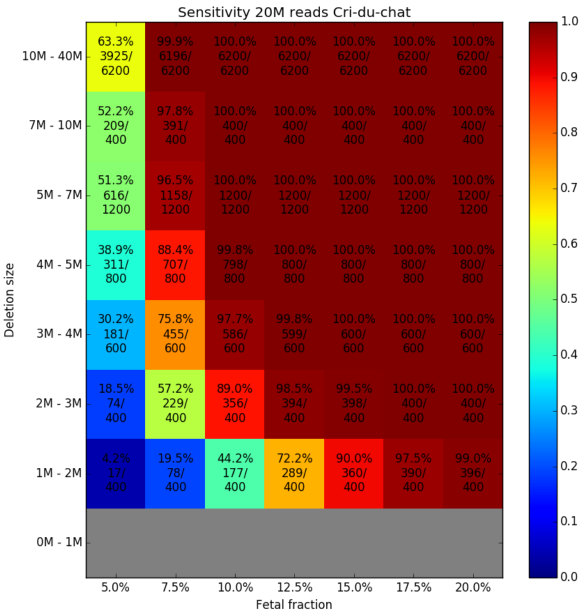
Sensitivity 20M reads Cri-du-chat
Deletion size
Fetal fraction
10M - 40M
63.3% 3925/ 6200
99.9% 6196/ 6200
100.0% 6200/ 6200
100.0% 6200/ 6200
100.0% 6200/ 6200
100.0% 6200/ 6200
100.0% 6200/ 6200
7M - 10M
52.2% 209/ 400
97.8% 391/ 400
100.0% 400/ 400
100.0% 400/ 400
100.0% 400/ 400
100.0% 400/ 400
100.0% 400/ 400
5M - 7M
51.3% 616/ 1200
96.5% 1158/ 1200
100.0% 1200/ 1200
100.0% 1200/ 1200
100.0% 1200/ 1200
100.0% 1200/ 1200
100.0% 1200/ 1200
4M - 5M
38.9% 311/ 800
88.4% 707/ 800
99.8% 798/ 800
100.0% 800/ 800
100.0% 800/ 800
100.0% 800/ 800
100.0% 800/ 800
3M - 4M
30.2% 181/ 600
75.8% 455/ 600
97.7% 586/ 600
99.8% 599/ 600
100.0% 600/ 600
100.0% 600/ 600
100.0% 600/ 600
2M - 3M
18.5% 74/ 400
57.2% 229/ 400
89.0% 356/ 400
98.5% 394/ 400
99.5% 398/ 400
100.0% 400/ 400
100.0% 400/ 400
1M - 2M
4.2% 17/ 400
19.5% 78/ 400
44.2% 177/ 400
72.2% 289/ 400
90.0% 360/ 400
97.5% 390/ 400
99.0% 396/ 400
0M - 1M
5.0% 7.5% 10.0% 12.5% 15.0% 17.5% 20.0%
0.0 0.1 0.2 0.3 0.4 0.5 0.6 0.7 0.8 0.9 1.0

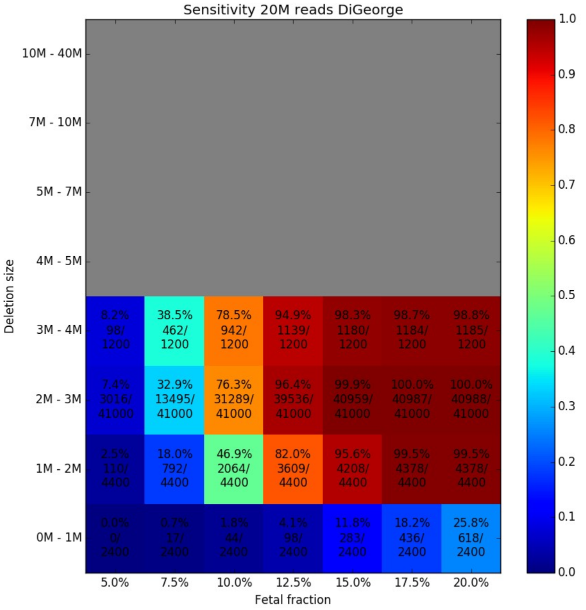


***Supplementary Figures 12.-16.:*** *Sensitivity of the prediction for different fetal fraction and microdeletion size for different syndromes. Read count was set to 20M in each sample. Detections 2M away from critical region are NOT reported.*

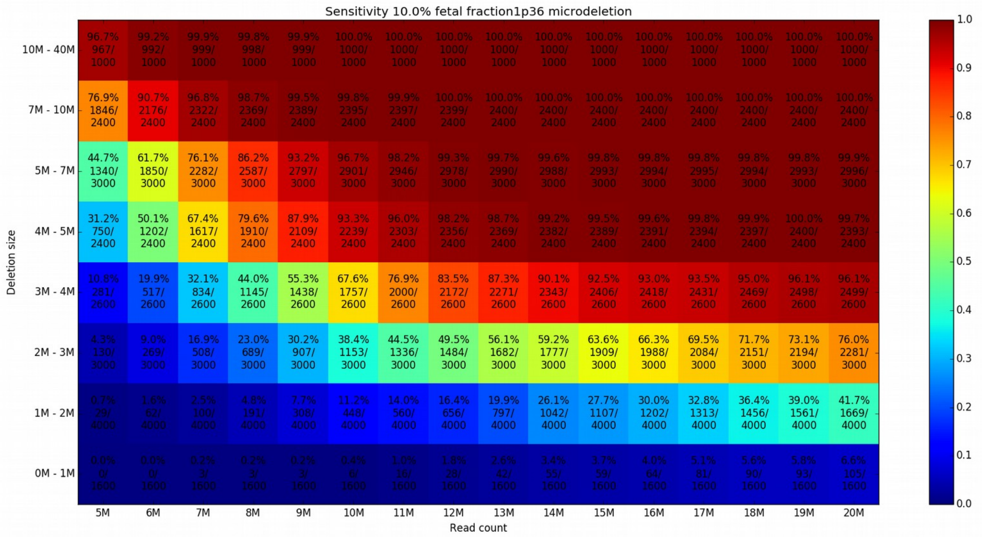

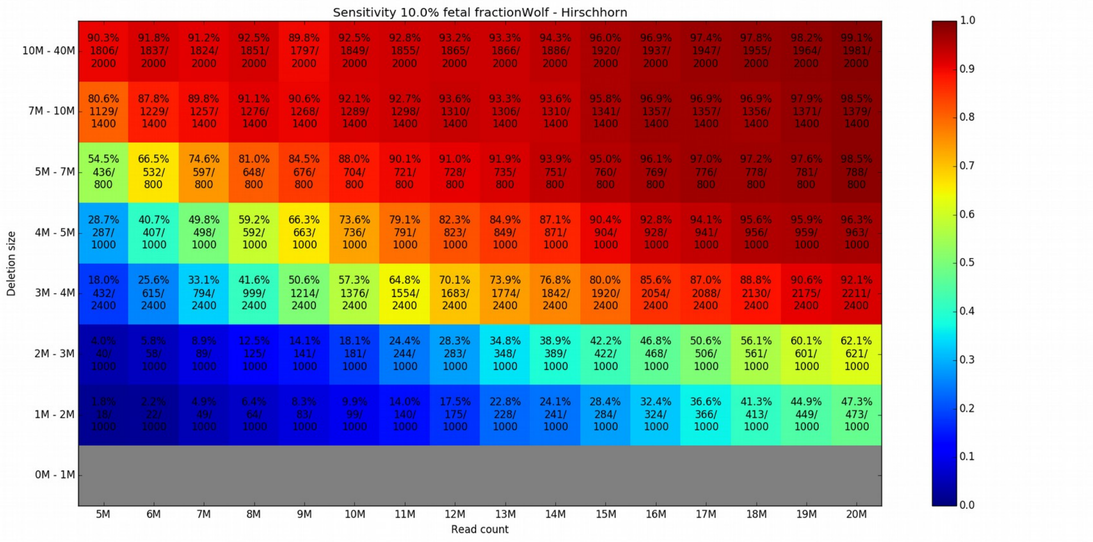
Sensitivity 10.0% fetal fractionWolf - Hirschhorn
Deletion size
Read count
5M 6M 7M 8M 9M 10M 11M 12M 13M 14M 15M 16M 17M 18M 19M 20M
10M - 40M: 90.3% 1806/ 2000; 91.8% 1837/ 2000; 91.2% 1824/ 2000; 92.5% 1851/ 2000; 89.8% 1797/ 2000; 92.5% 1849/ 2000; 92.8% 1855/ 2000; 93.2% 1865/ 2000; 93.3% 1866/ 2000; 94.3% 1886/ 2000; 96.0% 1920/ 2000; 96.9% 1937/ 2000; 97.4% 1947/ 2000; 97.8% 1955/ 2000; 98.2% 1964/ 2000; 99.1% 1981/ 2000
7M - 10M: 80.6% 1129/ 1400; 87.8% 1229/ 1400; 89.8% 1257/ 1400; 91.1% 1276/ 1400; 90.6% 1268/ 1400; 92.1% 1289/ 1400; 92.7% 1298/ 1400; 93.6% 1310/ 1400; 93.3% 1306/ 1400; 93.6% 1310/ 1400; 95.8% 1341/ 1400; 96.9% 1357/ 1400; 96.9% 1357/ 1400; 96.9% 1356/ 1400; 97.9% 1371/ 1400; 98.5% 1379/ 1400
5M - 7M: 54.5% 436/ 800; 66.5% 532/ 800; 74.6% 597/ 800; 81.0% 648/ 800; 84.5% 676/ 800; 88.0% 704/ 800; 90.1% 721/ 800; 91.0% 728/ 800; 91.9% 735/ 800; 93.9% 751/ 800; 95.0% 760/ 800; 96.1% 769/ 800; 97.0% 776/ 800; 97.2% 778/ 800; 97.6% 781/ 800; 98.5% 788/ 800
4M - 5M: 28.7% 287/ 1000; 40.7% 407/ 1000; 49.8% 498/ 1000; 59.2% 592/ 1000; 66.3% 663/ 1000; 73.6% 736/ 1000; 79.1% 791/ 1000; 82.3% 823/ 1000; 84.9% 849/ 1000; 87.1% 871/ 1000; 90.4% 904/ 1000; 92.8% 928/ 1000; 94.1% 941/ 1000; 95.6% 956/ 1000; 95.9% 959/ 1000; 96.3% 963/ 1000
3M - 4M: 18.0% 432/ 2400; 25.6% 615/ 2400; 33.1% 794/ 2400; 41.6% 999/ 2400; 50.6% 1214/ 2400; 57.3% 1376/ 2400; 64.8% 1554/ 2400; 70.1% 1683/ 2400; 73.9% 1774/ 2400; 76.8% 1842/ 2400; 80.0% 1920/ 2400; 85.6% 2054/ 2400; 87.0% 2088/ 2400; 88.8% 2130/ 2400; 90.6% 2175/ 2400; 92.1% 2211/ 2400
2M - 3M: 4.0% 40/ 1000; 5.8% 58/ 1000; 8.9% 89/ 1000; 12.5% 125/ 1000; 14.1% 141/ 1000; 18.1% 181/ 1000; 24.4% 244/ 1000; 28.3% 283/ 1000; 34.8% 348/ 1000; 38.9% 389/ 1000; 42.2% 422/ 1000; 46.8% 468/ 1000; 50.6% 506/ 1000; 56.1% 561/ 1000; 60.1% 601/ 1000; 62.1% 621/ 1000
1M - 2M: 1.8% 18/ 1000; 2.2% 22/ 1000; 4.9% 49/ 1000; 6.4% 64/ 1000; 8.3% 83/ 1000; 9.9% 99/ 1000; 14.0% 140/ 1000; 17.5% 175/ 1000; 22.8% 228/ 1000; 24.1% 241/ 1000; 28.4% 284/ 1000; 32.4% 324/ 1000; 36.6% 366/ 1000; 41.3% 413/ 1000; 44.9% 449/ 1000; 47.3% 473/ 1000
0M - 1M
0.0 0.1 0.2 0.3 0.4 0.5 0.6 0.7 0.8 0.9 1.0

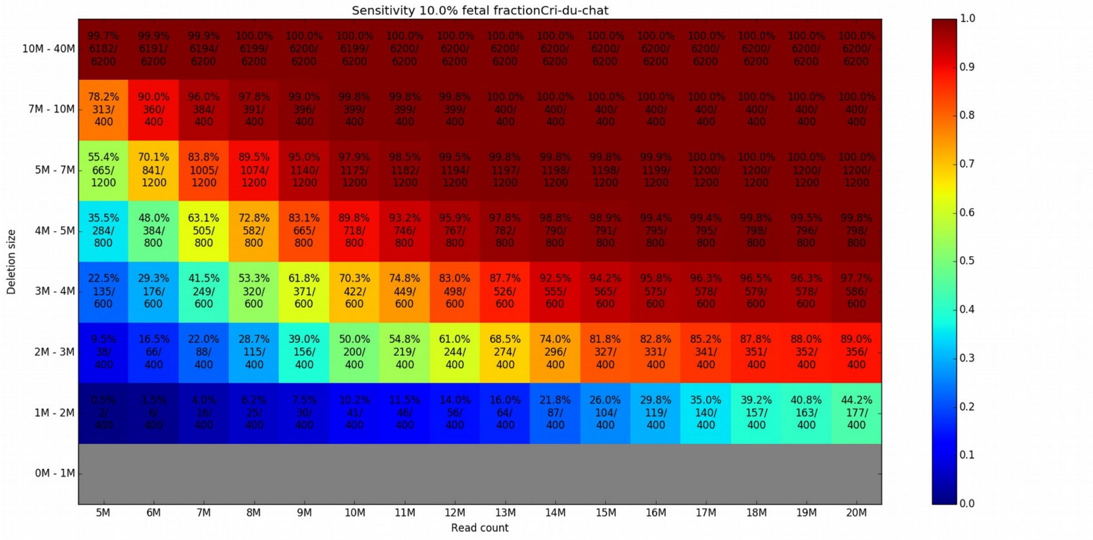
Sensitivity 10.0% fetal fractionCri-du-chat
Deletion size
Read count
5M 6M 7M 8M 9M 10M 11M 12M 13M 14M 15M 16M 17M 18M 19M 20M
10M - 40M: 99.7% 6182/ 6200; 99.9% 6191/ 6200; 99.9% 6194/ 6200; 100.0% 6199/ 6200; 100.0% 6200/ 6200; 100.0% 6199/ 6200; 100.0% 6200/ 6200; 100.0% 6200/ 6200; 100.0% 6200/ 6200; 100.0% 6200/ 6200; 100.0% 6200/ 6200; 100.0% 6200/ 6200; 100.0% 6200/ 6200; 100.0% 6200/ 6200; 100.0% 6200/ 6200; 100.0% 6200/ 6200
7M - 10M: 78.2% 313/ 400; 90.0% 360/ 400; 96.0% 384/ 400; 97.8% 391/ 400; 99.0% 396/ 400; 99.8% 399/ 400; 99.8% 399/ 400; 99.8% 399/ 400; 100.0% 400/ 400; 100.0% 400/ 400; 100.0% 400/ 400; 100.0% 400/ 400; 100.0% 400/ 400; 100.0% 400/ 400; 100.0% 400/ 400; 100.0% 400/ 400
5M - 7M: 55.4% 665/ 1200; 70.1% 841/ 1200; 83.8% 1005/ 1200; 89.5% 1074/ 1200; 95.0% 1140/ 1200; 97.9% 1175/ 1200; 98.5% 1182/ 1200; 99.5% 1194/ 1200; 99.8% 1197/ 1200; 99.8% 1198/ 1200; 99.8% 1198/ 1200; 99.9% 1199/ 1200; 100.0% 1200/ 1200; 100.0% 1200/ 1200; 100.0% 1200/ 1200; 100.0% 1200/ 1200
4M - 5M: 35.5% 284/ 800; 48.0% 384/ 800; 63.1% 505/ 800; 72.8% 582/ 800; 83.1% 665/ 800; 89.8% 718/ 800; 93.2% 746/ 800; 95.9% 767/ 800; 97.8% 782/ 800; 98.8% 790/ 800; 98.9% 791/ 800; 99.4% 795/ 800; 99.4% 795/ 800; 99.8% 798/ 800; 99.5% 796/ 800; 99.8% 798/ 800
3M - 4M: 22.5% 135/ 600; 29.3% 176/ 600; 41.5% 249/ 600; 53.3% 320/ 600; 61.8% 371/ 600; 70.3% 422/ 600; 74.8% 449/ 600; 83.0% 498/ 600; 87.7% 526/ 600; 92.5% 555/ 600; 94.2% 565/ 600; 95.8% 575/ 600; 96.3% 578/ 600; 96.5% 579/ 600; 96.3% 578/ 600; 97.7% 586/ 600
2M - 3M: 9.5% 38/ 400; 16.5% 66/ 400; 22.0% 88/ 400; 28.7% 115/ 400; 39.0% 156/ 400; 50.0% 200/ 400; 54.8% 219/ 400; 61.0% 244/ 400; 68.5% 274/ 400; 74.0% 296/ 400; 81.8% 327/ 400; 82.8% 331/ 400; 85.2% 341/ 400; 87.8% 351/ 400; 88.0% 352/ 400; 89.0% 356/ 400
1M - 2M: 0.5% 2/ 400; 1.5% 6/ 400; 4.0% 16/ 400; 6.2% 25/ 400; 7.5% 30/ 400; 10.2% 41/ 400; 11.5% 46/ 400; 14.0% 56/ 400; 16.0% 64/ 400; 21.8% 87/ 400; 26.0% 104/ 400; 29.8% 119/ 400; 35.0% 140/ 400; 39.2% 157/ 400; 40.8% 163/ 400; 44.2% 177/ 400
0M - 1M
0.0 0.1 0.2 0.3 0.4 0.5 0.6 0.7 0.8 0.9 1.0

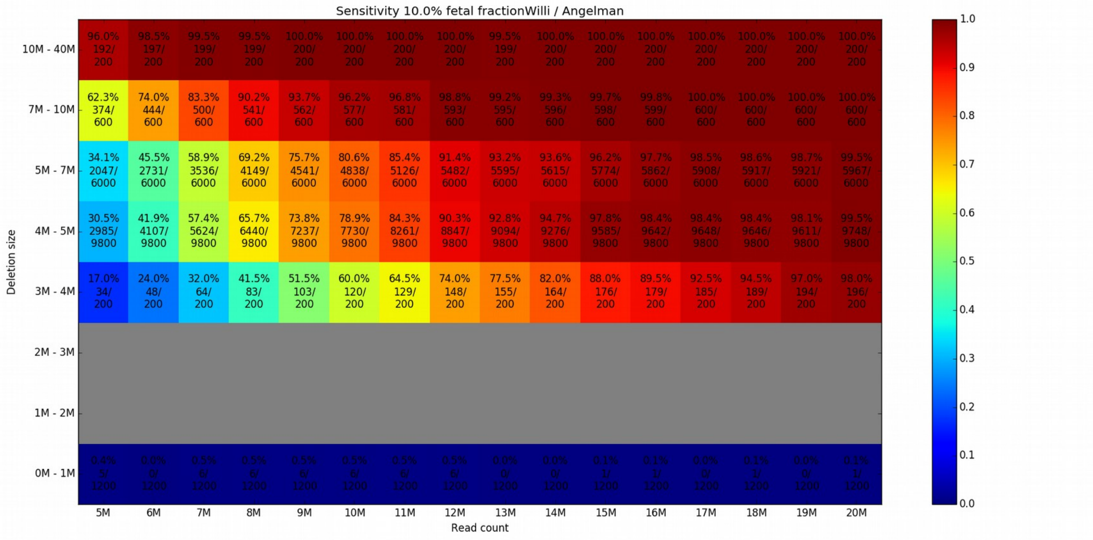
Sensitivity 10.0% fetal fractionWilli / Angelman
Deletion size
Read count
5M 6M 7M 8M 9M 10M 11M 12M 13M 14M 15M 16M 17M 18M 19M 20M
10M - 40M: 96.0% 192/ 200; 98.5% 197/ 200; 99.5% 199/ 200; 99.5% 199/ 200; 100.0% 200/ 200; 100.0% 200/ 200; 100.0% 200/ 200; 100.0% 200/ 200; 99.5% 199/ 200; 100.0% 200/ 200; 100.0% 200/ 200; 100.0% 200/ 200; 100.0% 200/ 200; 100.0% 200/ 200; 100.0% 200/ 200; 100.0% 200/ 200
7M - 10M: 62.3% 374/ 600; 74.0% 444/ 600; 83.3% 500/ 600; 90.2% 541/ 600; 93.7% 562/ 600; 96.2% 577/ 600; 96.8% 581/ 600; 98.8% 593/ 600; 99.2% 595/ 600; 99.3% 596/ 600; 99.7% 598/ 600; 99.8% 599/ 600; 100.0% 600/ 600; 100.0% 600/ 600; 100.0% 600/ 600; 100.0% 600/ 600
5M - 7M: 34.1% 2047/ 6000; 45.5% 2731/ 6000; 58.9% 3536/ 6000; 69.2% 4149/ 6000; 75.7% 4541/ 6000; 80.6% 4838/ 6000; 85.4% 5126/ 6000; 91.4% 5482/ 6000; 93.2% 5595/ 6000; 93.6% 5615/ 6000; 96.2% 5774/ 6000; 97.7% 5862/ 6000; 98.5% 5908/ 6000; 98.6% 5917/ 6000; 98.7% 5921/ 6000; 99.5% 5967/ 6000
4M - 5M: 30.5% 2985/ 9800; 41.9% 4107/ 9800; 57.4% 5624/ 9800; 65.7% 6440/ 9800; 73.8% 7237/ 9800; 78.9% 7730/ 9800; 84.3% 8261/ 9800; 90.3% 8847/ 9800; 92.8% 9094/ 9800; 94.7% 9276/ 9800; 97.8% 9585/ 9800; 98.4% 9642/ 9800; 98.4% 9648/ 9800; 98.4% 9646/ 9800; 98.1% 9611/ 9800; 99.5% 9748/ 9800
3M - 4M: 17.0% 34/ 200; 24.0% 48/ 200; 32.0% 64/ 200; 41.5% 83/ 200; 51.5% 103/ 200; 60.0% 120/ 200; 64.5% 129/ 200; 74.0% 148/ 200; 77.5% 155/ 200; 82.0% 164/ 200; 88.0% 176/ 200; 89.5% 179/ 200; 92.5% 185/ 200; 94.5% 189/ 200; 97.0% 194/ 200; 98.0% 196/ 200
2M - 3M
1M - 2M
0M - 1M: 0.4% 5/ 1200; 0.0% 0/ 1200; 0.5% 6/ 1200; 0.5% 6/ 1200; 0.5% 6/ 1200; 0.5% 6/ 1200; 0.5% 6/ 1200; 0.5% 6/ 1200; 0.0% 0/ 1200; 0.0% 0/ 1200; 0.1% 1/ 1200; 0.1% 1/ 1200; 0.0% 0/ 1200; 0.1% 1/ 1200; 0.0% 0/ 1200; 0.1% 1/ 1200
0.0 0.1 0.2 0.3 0.4 0.5 0.6 0.7 0.8 0.9 1.0

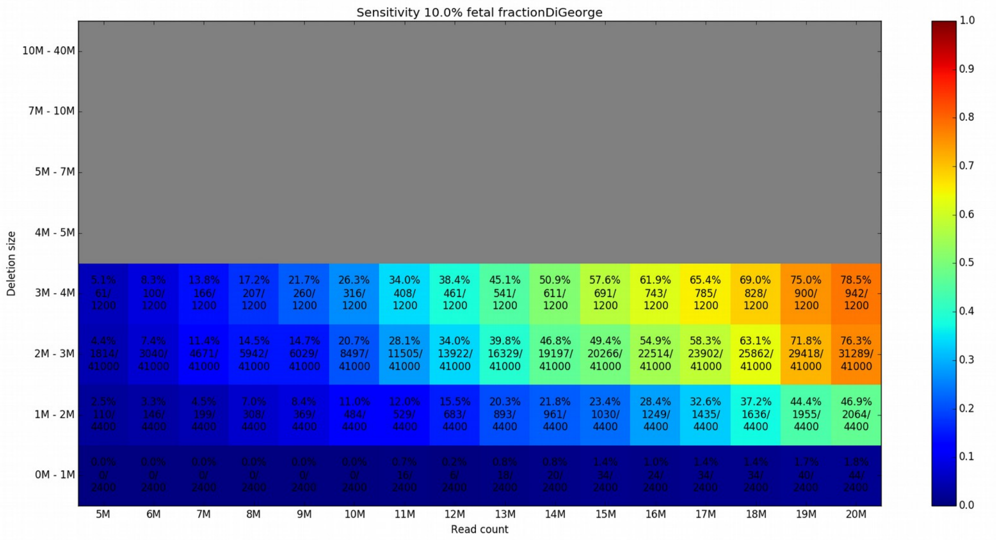


***Supplementary Figures 17.-21.:*** *Sensitivity of the prediction for different read count and microdeletion size. Fetal fraction was set to 10% for all samples. Detections 2M away from critical region are NOT reported.*

| name | read count | fetal fraction | detection | local z score | size | syndrome |
|---|---|---|---|---|---|---|
| NA17942 | 17.5M | 4.15% | ND | ND | 2.6M | DiGeorge |
| BA161216 | 25.27M | 4.50% | ND/grey zone | -2,48 | 3M | DiGeorge |
| NA22936 | 15.9M | 4.86% | D | -5,28 | 21M | 1p36 microdeletion |
| NA14124 | 16.47M | 5.11% | D | -8,35 | 17.7M | Cri-du-chat |
| BA34 | 19.24M | 5.90% | ND | ND | 0.9M | DiGeorge |
| NA09024 | 15.3M | 7.30% | D | -4,82 | 5.3M | Willi / Angelman |
| NA09024 | 12.66M | 7.60% | D | -5,79 | 5.3M | Willi / Angelman |
| NA17942 | 19.6M | 8.70% | D | -4,29 | 2.6M | DiGeorge |
| BA34 | 11M | 8.80% | ND | ND | 0.9M | DiGeorge |
| NA00072 | 8.3M | 9.10% | D | -16,98 | 9.3M | Wolf - Hirschhorn |
| NA22936 | 21.5M | 9.85% | D | -10,66 | 21M | 1p36 microdeletion |
| NA11515 | 19.31M | 10.60% | D | -5,59 | 6M | Willi / Angelman |
| BA34 | 9.9M | 10% | ND | ND | 0.9M | DiGeorge |
| BA161216 | 20.59M | 11.20% | D | -4,71 | 3M | DiGeorge |
| BA34 | 20.36M | 11.50% | D | -5,34 | 0.9M | DiGeorge |
| BA34 | 8.9M | 11.99% | D | -5,74 | 0.9M | DiGeorge |
| NA11515 | 16.79M | 12.20% | D | -9,02 | 6M | Willi / Angelman |
| NA09024 | 24.3M | 13.40% | D | -7,38 | 5.3M | Willi / Angelman |
| NA11515 | 15.71M | 13.90% | D | -9,54 | 6M | Willi / Angelman |
| NA00072 | 8.4M | 14.10% | D | -14,62 | 9.3M | Wolf - Hirschhorn |
| NA09024 | 15.37M | 14.50% | D | -10.25/-5.58 | 5.3M | Willi / Angelman |
| NA11515 | 19.26M | 14.60% | D | -7,72 | 6M | Willi / Angelman |
| NA00072 | 16.2M | 16.40% | D | -14,3 | 9.3M | Wolf - Hirschhorn |
| NA17942 | 14.5M | 16.69% | D | -7,11 | 2.6M | DiGeorge |
| BA34 | 21.52M | 17.30% | D | -6,52 | 0.9M | DiGeorge |
| NA17942 | 19.45M | 17.30% | D | -6,71 | 2.6M | DiGeorge |
| BA161216 | 20.39M | 20.10% | D | -8,27 | 3M | DiGeorge |
| NA17942 | 20.39M | 20.10% | D | -8,27 | 2.6M | DiGeorge |
| NA09024 | 16.1M | 20.50% | D | -13.50/-5.71 | 5.3M | Willi / Angelman |

***Supplementary Table 1.:*** *Detailed description of the control samples and their detection accuracy.*

| Identification of a syndrome | | Number of pathogenic deletions | | Mean size of pathogenic deletions | |
|---|---|---|---|---|---|
| **Band** | **Name** | **ISCA** | **DECIPHER** | **ISCA** | **DECIPHER** |
| 1p36 | 1p36 deletion | 102 (100) | 54 (52) | 4,131,271 | 3,174,263 |
| 4p16.3 | Wolf-Hirschhorn | 48 (48) | 27 (27) | 6,713,778 | 4,819,991 |
| 5p15 | Cri-du-chat | 50 (50) | 24 (24) | 14,887,247 | 8,873,724 |
| 15q11 | Angelman / Prader-Willi | 90 (90) | 133 (133) | 5,061,279 | 2,534,265 |
| 22q11.21 | DiGeorge | 280 (245) | 136 (120) | 2,368,692 | 2,444,038 |

| Critical regions | | Pathogenic deletions from DECIPHER not in ISCA critical region |
|---|---|---|
| **ISCA** | **DECIPHER** | |
| chr1: 564,424 - 21,598,492 | chr1: 120,840 - 22,542,549 | 3 |
| chr4: 85,040 - 2,010,761 | chr4: 71,552 - 5,506,588 | 2 |
| chr5: 1 - 15,678,560 | chr5: 113,576 - 18,992,827 | 2 |
| chr15: 22,779,922 - 28,559,437 | chr15: 22,373,311 - 28,560,803 | 2 |
| chr22: 18,661,724 - 21,505,417 | chr22: 18,890,162 - 21,464,119 | 15 |

***Supplementary Table 2.:*** *Number, mean size of pathogenic regions and resulting critical regions as reported by our study and DECIPHER database. The number of pathogenic regions overlapping its corresponding critical region is reported in brackets. Finally, the number of pathogenic deletions from DECIPHER database that does not overlap ISCA database is low.*
